# Identification of limb-specific *Lmx1b* auto-regulatory modules with Nail-Patella Syndrome pathogenicity

**DOI:** 10.1101/2020.08.10.243915

**Authors:** Endika Haro, Florence Petit, Charmaine U. Pira, Conor D. Spady, Lauren A. Ivey, Austin L. Gray, Fabienne Escande, Anne-Sophie Jourdain, Andy Nguyen, Florence Fellmann, Jean-Marc Good, Christine Francannet, Sylvie Manouvrier-Hanu, Marian A. Ros, Kerby C. Oberg

**Affiliations:** Department of Pathology and Human Anatomy, Loma Linda University School of Medicine, Loma Linda, CA USA; Instituto de Biomedicina y Biotecnología de Cantabria, CSIC–SODERCAN-Universidad de Cantabria, Santander, Spain; Clinique de Génétique, CHU Lille, F-59000 Lille, France; EA7364 RADEME, Université de Lille, F-59000 Lille, France; Laboratoire de Biochimie et Biologie Moléculaire, CHU Lille, F-59000 Lille, France; Service de Médecine Génétique, Centre Hospitalier Universitaire Vaudois, Lausanne, Switzerland; Service de génétique médicale, CHU Estaing, Clermont-Ferrand, France

**Keywords:** Limb patterning, limb dorsalization, autoregulation, human, nail-patella syndrome, murine, *cis*-regulatory modules, Lmx1b

## Abstract

*LMX1B* haploinsufficiency causes Nail-patella syndrome (NPS; MIM 161200), characterized by nail dysplasia, absent/hypoplastic patellae, chronic kidney disease, and glaucoma. Accordingly, in mice *Lmx1b* has been shown to play crucial roles in the development of the limb, kidney and eye. Although one functional allele of murine *Lmx1b* appears adequate for development, *Lmx1b* null mice display ventral-ventral distal limbs with abnormal kidney, eye and cerebellar development, more disruptive, but fully concordant with NPS. Interestingly, in *Lmx1b* functional knockouts (KOs), *Lmx1b* transcription in the limb is decreased nearly 6-fold indicating autoregulation. Herein, we report on two conserved *<u>L</u>mx1b*-<u>a</u>ssociated *cis*-<u>r</u>egulatory <u>m</u>odules (*LARM1* and *LARM2)* that are bound by Lmx1b, amplify *Lmx1b* expression in the limb and are necessary for Lmx1b-mediated limb dorsalization. Remarkably, we also report on two NPS patient families with normal *LMX1B* coding sequence, but loss-of-function variations in the *LARM1/2* region, stressing the role of regulatory modules in disease pathogenesis.

## Introduction

The LIM homeodomain transcription factor Lmx1b is responsible for limb dorsalization. In the limb, Lmx1b is induced by Wnt7a from the dorsal ectoderm and its expression is restricted to the dorsal mesoderm^1,2^. Loss of Lmx1b function in mice results in loss of dorsal autopod (hand/foot) and zeugopod (forearm/leg) patterning; the autopods have a symmetrical ventral-ventral phenotype with dorsal footpads, loss of dorsal hair follicles, absence of nails, and a symmetrical ventral-ventral pattern of muscles, tendons and ligaments. Besides the limb, mice lacking functional *Lmx1b* exhibit abnormal eye, cerebellar, and kidney development which accounts for the perinatal lethality^3^. In contrast to mice, single allele variations in humans that disrupt LMX1B function cause Nail-Patella Syndrome (NPS; MIM 161200)^4,5^. This autosomal dominant condition is characterized by nail dysplasia, absent or hypoplastic patellae, bone fragility, premature osteoarthritis, chronic kidney disease, and ocular anomalies. The human syndromic phenotype is less dramatic than exhibited in homozygous KO mice implying that haploinsufficiency or an inadequate level of LMX1B is responsible for the syndromic features.

In the absence of Lmx1b function, transcription of the *Lmx1b* mRNA is decreased nearly 6-fold in developing mouse limbs suggesting that one function of Lmx1b is the auto-amplification of its own expression^6^. Lmx1b-targeted chromatin immunoprecipitation combined with high-throughput sequencing (ChIP-seq) during limb dorsalization identified two highly conserved Lmx1b-bound cis-regulatory modules (CRMs) 60 kb upstream of the *Lmx1b* gene^7^. CRMs are DNA sequences enriched in transcription factor binding sites that regulate associated genes in a time and tissue specific manner. Lmx1b-binding to CRMs upstream of its own coding sequence provide a mechanism by which Lmx1b could auto-amplify its own expression.

In this study we have characterized and functionally validated these two <u>L</u>mx1b <u>a</u>ssociated <u>r</u>egulatory <u>m</u>odules, that we term *LARM1* and *LARM2*. We show that the activity of *LARM1*/2 overlaps the expression pattern of *Lmx1b* in the dorsal limb mesoderm and is conserved across vertebrates including humans. Removal of *LARM1*/2 with CRISPR-Cas9, resulted in a limb-phenotype identical to that of animals lacking functional *Lmx1b* with marked reduction in *Lmx1b* expression and loss of limb dorso-ventral asymmetry, but without any other Lmx1b-related organ system affected. These data establish *LARM1*/2 as limb-specific *Lmx1b* enhancers necessary for amplifying the level of Lmx1b expression in the limbs. Interestingly, about 10% of patients with the NPS phenotype lack a variation in the *LMX1B* coding sequence^8^. We found two NPS patient families lacking coding sequence variations that had *LARM* variations that disrupt human LARM activity highlighting the important role of regulatory modules in disease pathogenesis.

## Results

### *Lmx1b*-associated regulatory modules are active during limb dorsalization, require Lmx1b binding sites for activity, and are activated by LMX1B

We recently identified two Lmx1b-bound CRMs, *LARM1* and *LARM2,* 60 and 66 kb upstream of the mouse *Lmx1b* gene, respectively (Fig. 1A)^7^, that are associated with active chromatin marks (p300, H3K27ac, H3K4me2, RNAPOL2 and Med12) during limb development (Fig. 1B). Interestingly, *LARM*2, the more distant module, also associates with inhibitory marks (H3K27me3). For a given cell, histone acetylation (H3K27ac) and methylation (H3K27me3) are mutually exclusive; their coexistence indicates tissue heterogeneity. This pattern is consistent with *LARM2* being only accessible and active in the dorsal limb compartment^9^.

**Figure 1.**
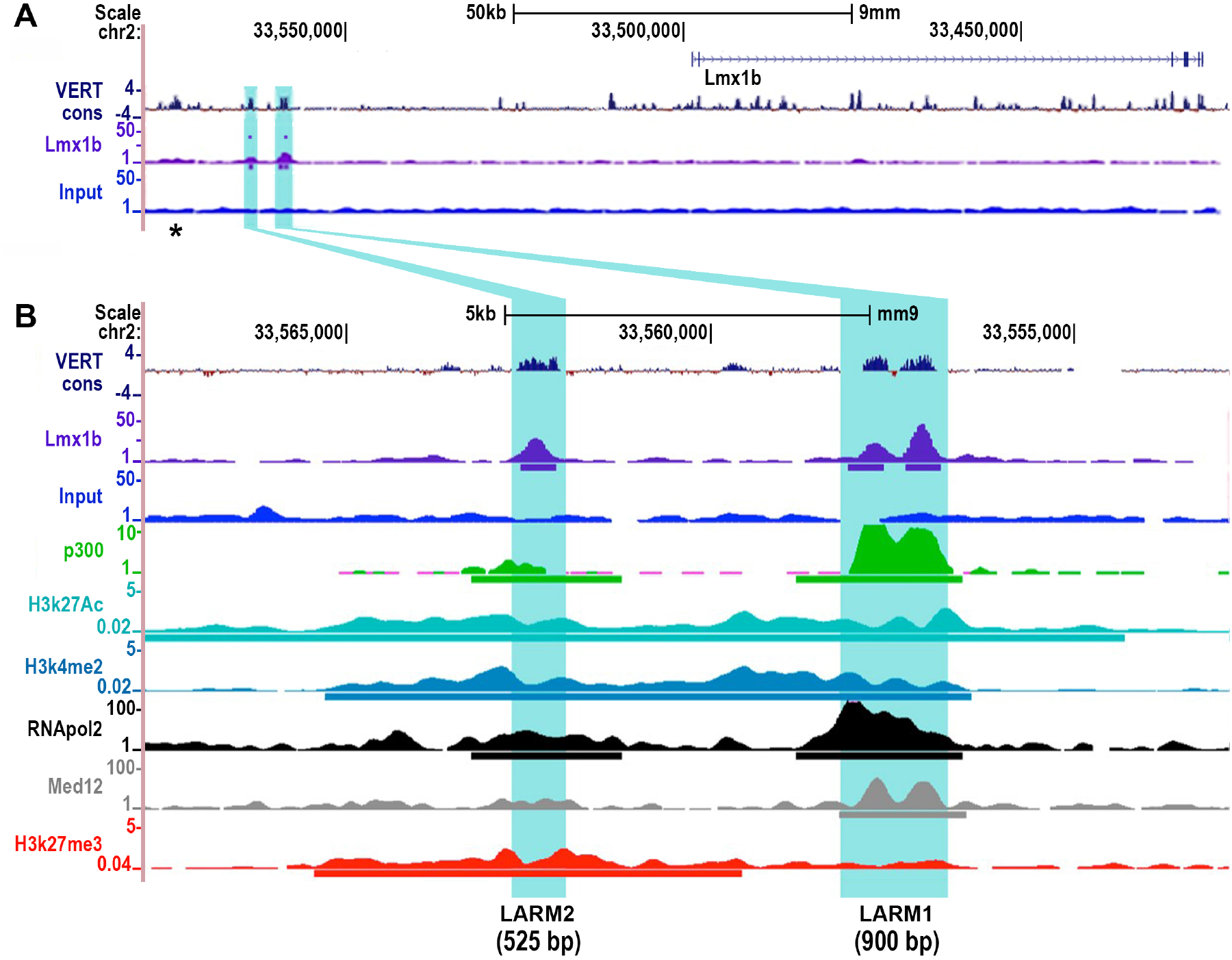
*LARM1* and *LARM2* are conserved, bound by Lmx1b, and associated with active chromatin marks. A) UCSC genome screenshot displaying the *Lmx1b* locus and the associated *cis*-regulatory modules *LARM1* and *LARM2* (highlighted in blue) showing vertebrate conservation (VERT cons), Lmx1b binding (Lmx1b-targeted ChIP-seq)^7^ and input^7^. B) Magnification of the putative enhancer region displaying the overlap with active enhancer associated regulatory marks present in limb buds. From top to bottom: vertebrate conservation (VERT cons), ChIP-seq tracks for Lmx1b^7^, control Input limb DNA^7^, p300^20^, histone 3 acetylation at lysine 27 (H3k27Ac)^21^, histone 3 dimethylation at lysine 4 (H3k4me2)^22^, RNA polymerase II^23^, Med 12^23^ and histone 3 trimethylation at lysine 27 (H3Kk7me3)^22^. Note that LARM1 consists of two conserved peaks, both of which are recognized by Lmx1b-targeted ChIP-seq^7^.

Using GFP reporter constructs in a chicken electroporation bioassay^10,11^, we found that both *LARM1* and *LARM*2 demonstrate enhancer activity within the dorsal limb mesoderm coincident with the *Lmx1b* expression domain (Fig. 2A, 2F). Conservation analysis using the VISTA browser (Fig. 1B) subdivided *LARM1* into two conserved regions, a 5’ element containing one conserved potential Lmx1b binding site and a 3’ element with three conserved binding sites (based on the reported TMATWA binding motif)^7^ (Fig. 2A). Surprisingly, the isolated 5’ *LARM1* element did not show reporter activity, while the isolated 3’ *LARM1* element showed strong activity in the limb mesoderm but with no dorsal bias (Fig. 2C). Interestingly, the restriction of *LARM1* activity to the dorsal mesoderm requires the Lmx1b binding site within the 5’ element as site-directed mutagenesis expanded enhancer activity into the ventral mesoderm (Fig. 2D). In contrast, mutation of any of the 3 predicted Lmx1b binding sites in the 3’ element resulted in markedly reduced, but still dorsally restricted LARM1 activity (Fig. 2D-E). These findings are counterintuitive since Lmx1b is only expressed in the dorsal limb mesoderm. A reasonable interpretation is that the putative Lmx1b binding site within the 5’ *LARM1* element (TTATTA) can bind with other transcription factors that silence the 3’ *LARM1* enhancer activity or promote chromatin conformation that limits enhancer-promoter interaction. In the dorsal mesoderm, Lmx1b would compete for this binding site, repress the 3’ *LARM1* element, and drive enhancer activity. Collectively, our data implies that *LARM1* is composed of a 3’ enhancer (*LARM1*e) and a 5’ silencer (*LARM1*s) that blocks ventral activity, thereby restricting its function to the dorsal limb.

**Figure 2.**
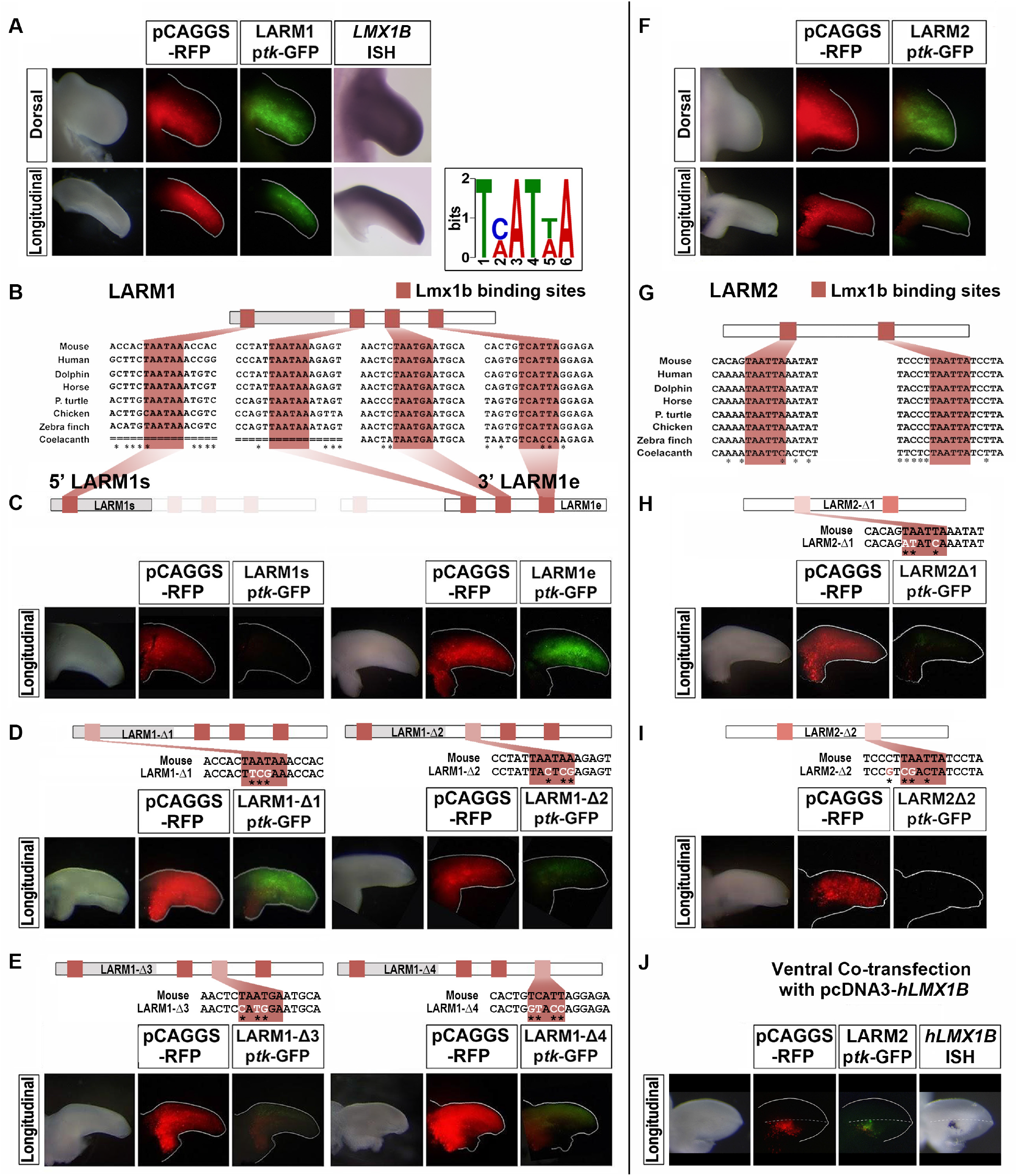
The Lmx1b binding sites are necessary for dorsal *LARM1* and *LARM2* activity. Activity of *LARM1* (A-E) and *LARM2* (F-J) in chick wing buds 48 hrs after electroporation of the respective *LARM*-reporter construct. For each assay/experiment a bright field view of the electroporated limb bud is accompanied by a fluorescence image of RFP (red) demonstrating transfection efficiency and a fluorescent image of GFP (green) demonstrating enhancer activity. Dorsal or longitudinal views as indicated. The longitudinal views illustrate activity along the dorsal-ventral axis (dorsal on top) A) *LARM1* has activity restricted to dorsal mesoderm (n=22) coincident with *LMX1B* expression (*LMX1B* ISH shown for comparison). Inset showing the TMATWA consensus DNA binding motif for Lmx1b. B) Conserved Lmx1b binding sites are shaded clay-red in *LARM1* schematics and sequences. Sequence variations across species is indicated as an asterisk below the alignment. C) Analysis of the isolated *3’LARM1* element (*LARM1s*) does not convey enhancer activity (n=4), while the isolated *5’LARM1* element (*LARM1e*) is active throughout the limb bud (n=16), in both dorsal and ventral mesoderm. D) Left panel: Interestingly, site-directed mutagenesis of the 5’ *LARM1s* Lmx1b binding site in the full *LARM1* construct (*LARM1-Δ1*, n=5) expands the activity to both dorsal and ventral mesoderm indicating that the binding site is necessary for the restriction of dorsal activity. D) right panel and E) Disruption of any of the Lmx1b binding sites present in the 3′ *LARM1e* element leads to a marked reduction in enhancer activity (LARM1-Δ2, n=7; LARM1-Δ3, n=5; LARM1-Δ4, n=5). F) *LARM2* enhancer activity is restricted to dorsal mesoderm (n=13) coincident with *LMX1B* expression (shown in A). G) Two highly conserved Lmx1b binding sites are present in *LARM*2 (shaded clay-red as in B). H and I) Disruption of either binding site leads to a loss in *LARM*2 activity (LARM2-Δ1, n=5; LARM2-Δ2, n=6). J) Ectopic expression of human LMX1B in the ventral mesoderm is able to drive activity of the co-transfected *LARM2*-reporter construct (n=3). The human *LMX1B* probe illustrates the ectopic human *LMX1B* expression by *in situ* hybridization, but does not cross-react with chicken *LMX1B* to display its dorsal expression. Nucleotides altered by site-directed mutagenesis are indicated in white and by an asterisk below the sequence.

In contrast, *LARM*2 is composed of a single conserved element containing two highly conserved predicted Lmx1b binding sites (Fig. 2G). The dorsally restricted activity of *LARM*2 is abolished by disruption of either binding site (Fig. 2H, I) suggesting that *LARM2* is a positive *Lmx1b* enhancer whose activity depends on Lmx1b. Furthermore, the observation that *LARM*2 reporter activity is activated in ventral limb mesoderm by ectopically expressing human LMX1B fully corroborates this conclusion (Fig. 2J). These results, together with the pattern of chromatin marks in the *LARM2* enhancer (Fig. 1B), suggest that Lmx1b may also play a role in chromatin activation within the dorsal compartment.

Finally, evaluation of previously published capture C experiments^12^ shows that *LARM1/2* physically interact with the *Lmx1b* promoter (Fig. S1). Thus, the observation that both *LARM1* and *LARM2* are bound by Lmx1b^7^, display dorsal restricted activity in limb buds, require Lmx1b binding sites for activity and interact with the Lmx1b promoter, in addition to human LMX1B being sufficient to activate *LARM2* in the ventral mesoderm, collectively support the concept that *LARM1*/2 are bona fide Lmx1b autoregulatory enhancers. Interestingly, one of the enhancers of *Apterous* (*ap*), the *Drosophila* homologue of *Lmx1b*, is maintained by an autoregulatory loop, albeit indirectly through the ap targets vestigial and scalloped (Vg/Sd)^13^.

### *LARM* activity is required for limb-specific *Lmx1b* amplification and distal dorsal limb patterning

To determine their functional role in *Lmx1b* regulation, we deleted *LARM1/2* by CRISPR-Cas9. Mice homozygous for a 7.6 kb deletion encompassing both *LARM1* and *LARM2* exhibited a limb phenotype similar to that observed in the absence of functional Lmx1b^3^ consisting in loss of dorsalization with distal ventral-ventral limbs that involved the skeleton, muscles and integument (Fig. 3). Micro computed tomography (microCT) demonstrated biventral distal skeletal elements (Fig. 3B-B’) with dorso-ventral symmetrical distal phalanges (Fig. 3C-C’), sesamoid bones (Fig. 3D-D’) and tali (Fig. 3E-E’). In addition, the patella, the dorsal most structure of the knee, is absent (also a notable feature in *Lmx1b* KO mice and NPS patients) (Fig. 3F-F’). These skeletal abnormalities were accompanied by corresponding muscular abnormalities (Fig. 3G-G’).

**Figure 3.**
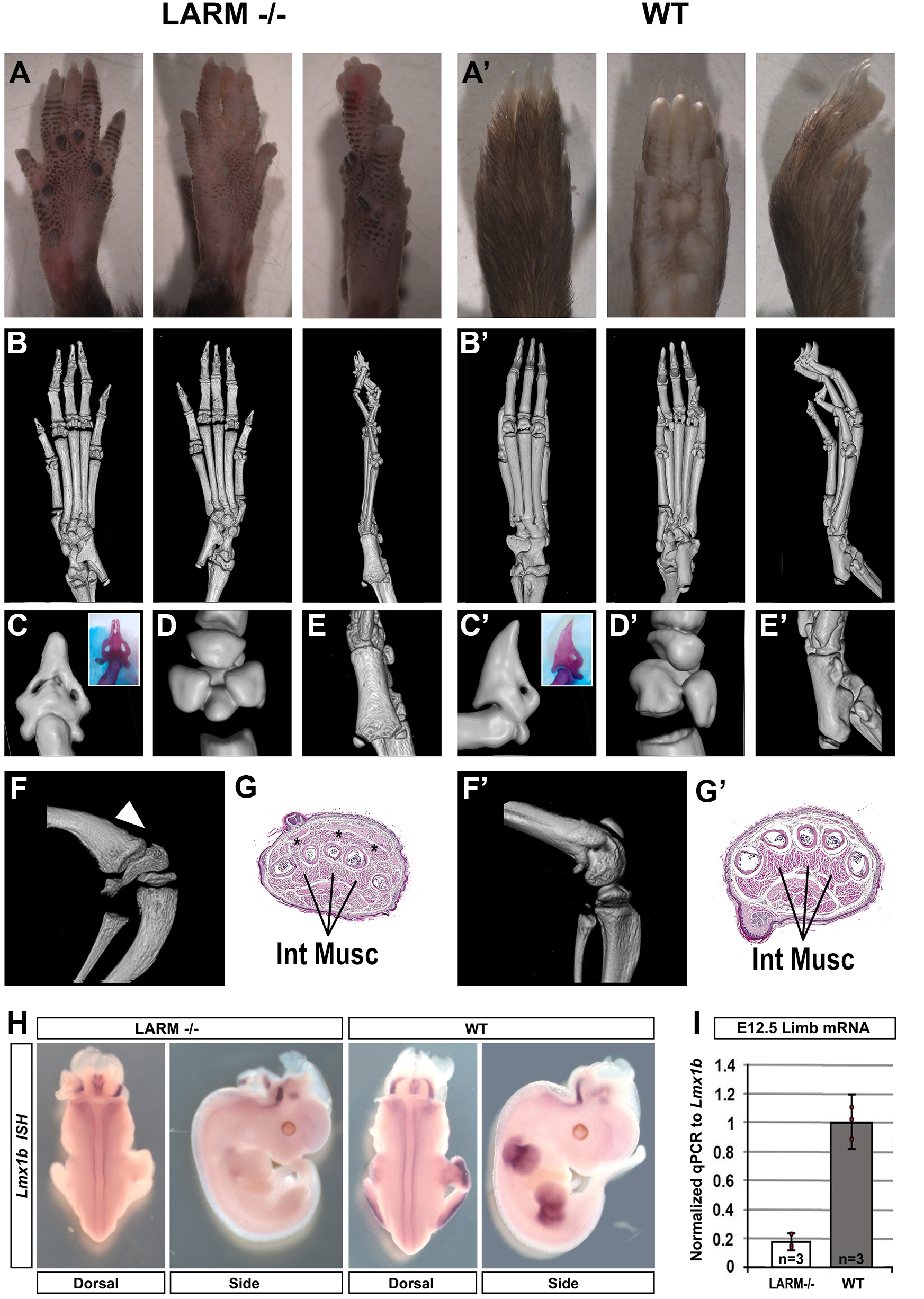
Mice deficient of *LARM1/2* exhibit a ventral-ventral limb phenotype. A-A’) Dorsal, ventral and lateral gross morphology and; B-B’) microCT scan views of a three-week-old *LARM1/2* homozygous mouse showing footpad development and the absence of nails and hair in the dorsal autopod. C-E and C′-E′) Magnified views of the digit tips, the sesamoid bones, and tali display dorsoventral symmetry in the absence of *LARM1/2*. F-F’) The patella is absent in *LARM1/2* KOs (hindlimb lateral view, white arrowheads). G-G′) Transverse sections of the autopod show duplicated flexor tendons and intrinsic muscles (Int M) (★) in the *LARM1/2* KO. *In situ* hybridization of *Lmx1b* expression in limb buds is below detection in animals lacking the Lmx1b associated regulatory modules (*LARM1/2*), while expression in the neural tube is equivalent. I) Quantitative PCR of *Lmx1b* mRNA demonstrates a marked reduction in *LARM1/2* KO mice ~ 20% of normal (WT) mice. \*\*\**P*<0.01 (two-tailed, unpaired *t*-test, *n*=biological replicates, each replicate is depicted as a red dot).

*Lmx1b* KO mice die shortly after birth, however, *LARM1/2* mutants are viable suggesting a more restricted phenotype. In *Lmx1b* KO mice the development of the calvaria, cerebellum, kidney and eye, are affected^3^, but these organs appear normal in the absence of *LARM1/2* indicating the limb-specific function of *LARM1/2* (Fig. S2). The analysis of *Lmx1b* expression in *LARM1/2* KO embryos by whole mount *in situ* hybridization showed a normal pattern except in the limb, where it was below detection limits (Fig. 3H). Analysis of limb RNA by qPCR at e12.5 demonstrated more than a 5-fold higher level of *Lmx1b* expression in normal mice compared to mice lacking the *LARM* region (Fig. 3I) similar to what has been reported in *Lmx1b* functional KO mice^6^. Collectively, our results establish *LARM1/2* as limb-specific *Lmx1b* CRMs that are absolutely necessary for normal *Lmx1b* transcription levels and dependent on Lmx1b for activity.

### Human *LARM1/2* activity and role in Nail-Patella Syndrome

*LARM1/2* are conserved in Humans including the LMX1B binding sites (Fig. 1 and 2). We isolated the *hLARM* sequences and demonstrated dorsally restricted activity in the chick bioassay, either isolated or together (Fig. 4A). We also evaluated the *hLARM* sequences in transgenic mice at e12.5 (Fig. 4B). The *LARM* transgenes displayed a limb-restricted and dorsally accentuated activity. *LARM1* exhibited accentuated activity in the dorsal limb mesoderm, but weaker activity was also evident in the distal ventral aspect. *LARM2* was tightly restricted to the dorsal mesoderm but also lacked activity in the fifth digital ray (Fig. 4B, arrow). The transgene including the entire *LARM* region had dorsally restricted expression with the exception of a small ventral patch in the presumptive carpal/tarsal region (Fig. 4B, arrowhead), suggesting a cooperative implementation of *LARM1* and *LARM2* activities for the refinement of dorsal restriction/enhancement.

**Figure 4.**
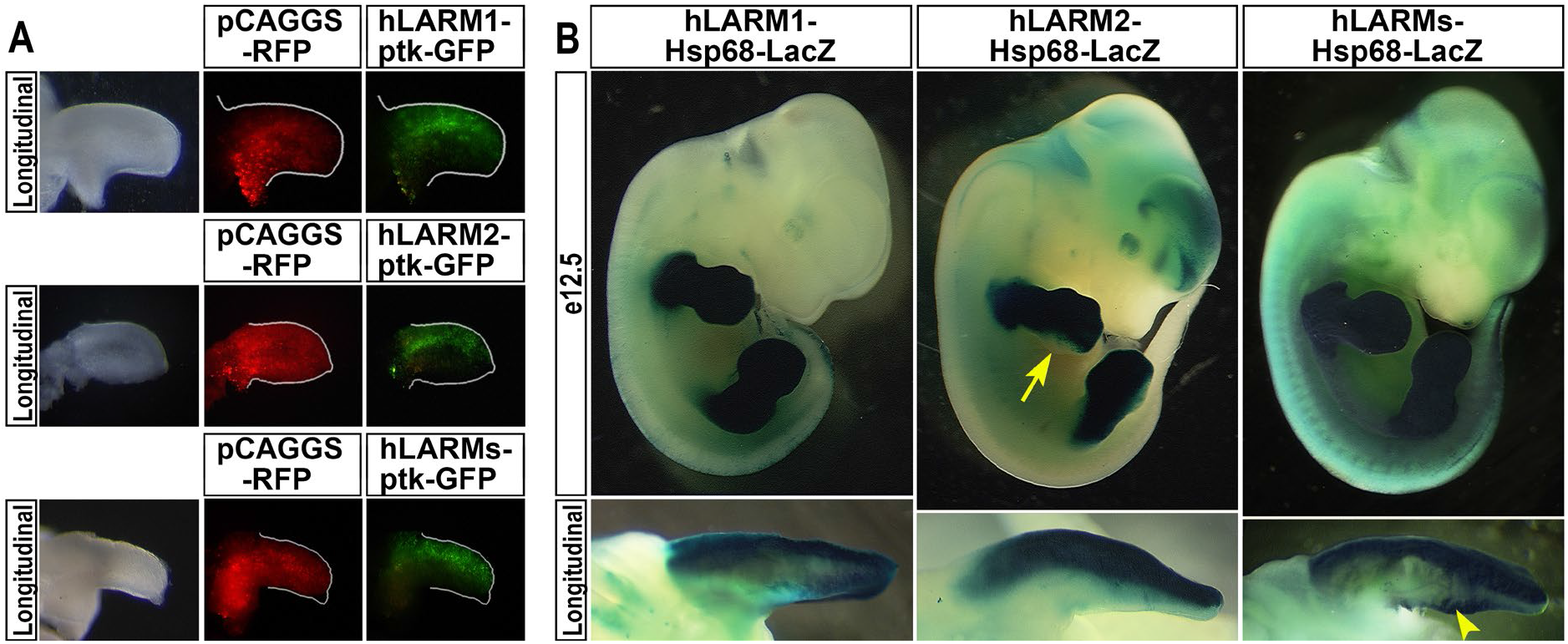
The human *LARM* region also exhibits dorsal enhancer activity. A) Human *LARM* constructs electroporated into chick wing buds display dorsally restricted expression (*hLARM1*, n=13; *hLARM2*, n=7; *hLARM1/2* (*hLARMs*), n=3). B) Similarly, transgenic mice containing the human *LARM* sequences linked to a LacZ reporter demonstrate dorsally accentuated activity. All three *LARM1* transgenic embryos displayed limb-restricted, dorsally accentuated activity. *LARM2* transgenic embryos exhibited tight dorsally restricted activity in the limb (5/5). Staining was reduced-to-absent in the posterior distal autopod mesoderm (5^th^ digit region, yellow arrow). Transgenic embryos containing both *hLARM1/2* (~8 kb, *hLARMs*) also showed activity restricted to the dorsal limb (7/7). However, focal ventral activity at the zeugopod/autopod junction was also evident (yellow arrowhead).

We also explored the *LARM* region in 11 unrelated patients affected with NPS lacking sequence or copy number variation of the *LMX1B* coding region. Five of these patients were reported in a recent study^8^. In one proband (IV-7), we identified a 4.5 kb heterozygous deletion encompassing a conserved region adjacent to *LARM1* and all of *LARM2* (Family 53 from Ghoumid et al.^8^) (Fig. 5A, B). The *LARM* deletion was shown to segregate in one affected first cousin (IV-1) and inferred from that result that two other affected individuals are obligate carriers (III-1 and III-4). Remarkably, individuals from this family exhibited nail dysplasia and patella hypoplasia, without ocular or renal involvement (Fig. 5C-N individual IV-7 and Fig. 5H-J individual IV-1).

**Figure 5.**
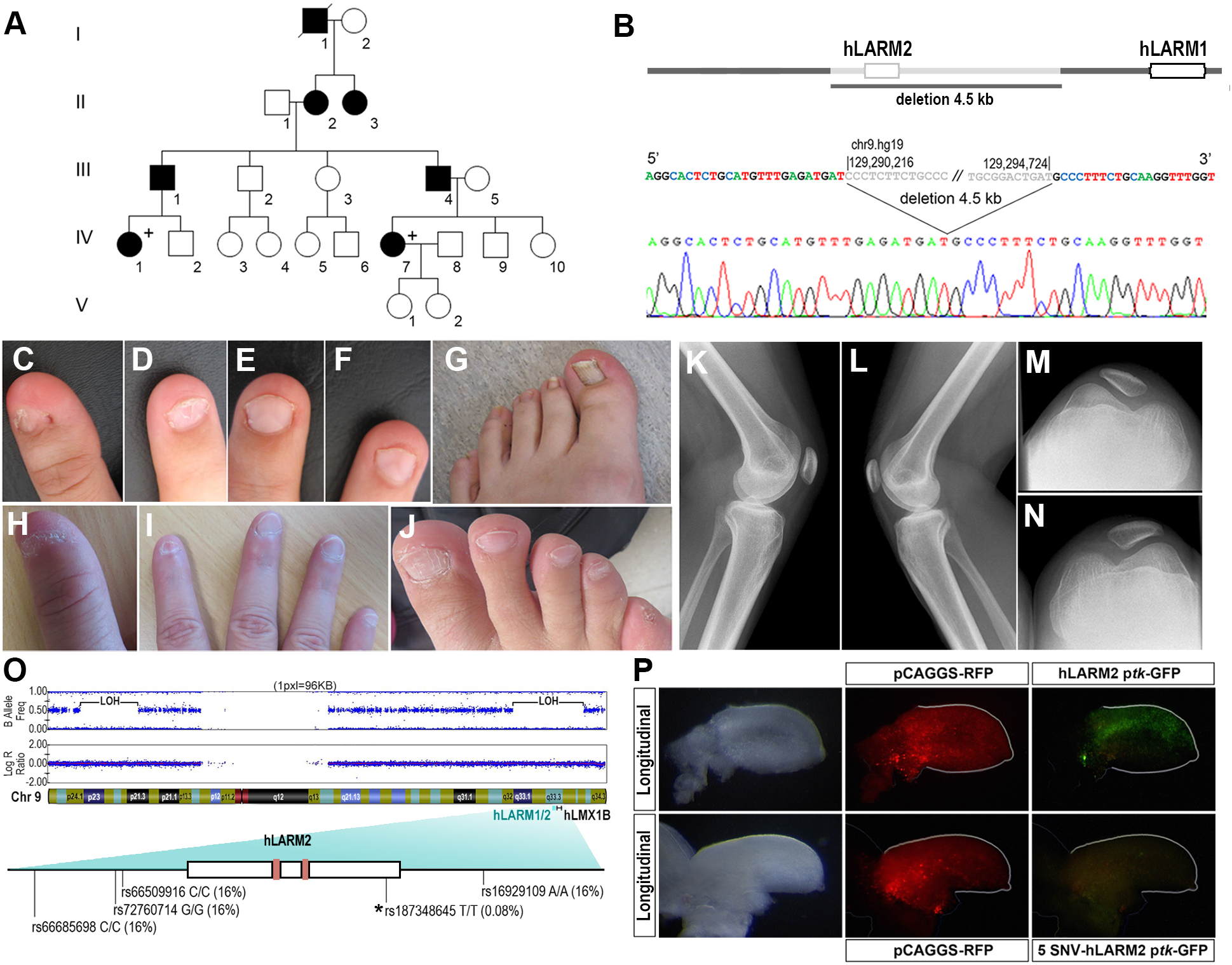
Clinical features of *LARM* loss-of-function. A) Pedigree of a family with a LARM deletion. B) The 4.5 kb region deleted removes all of *LARM2*. C-G and K-N) Phenotypic images of individual IV-7. C, D) Koïlonychia of thumb and 2^nd^ finger. E,F) Triangular lunulae of 3^rd^ and 4^th^ fingers. G) Nail dysplasia of the hallux showing longitudinal striations. H-J) Phenotypic images of individual IV-1. H) Koïlonychia of thumb. I) Hypoplastic nails, ungueal dysplasia of 2^nd^ finger. J) Ungueal dysplasia of right foot predominating on 1^st^ and 5^th^ toes. K-N) Knee x-rays showing bilateral hypoplasia of the patella. O) Schematic of the patient’s chromosome 9 showing large segments of the chromosome with loss of heterozygosity (LOH), i.e. homozygosity, when comparing the allele frequencies to the Log R ratio of the alleles. One of the homozygous regions includes the LARM-LMX1B locus. The homozygous *hLARM2* sequence showing the five single nucleotide variations (SNVs). The asterisk (★) indicates the rare (0.08%) sequence in *LARM2*. The LMX1B binding sites are indicated as clay-red boxes. P) Using site-directed mutagenesis, we generated a human *LARM2* construct containing the patient′s 5 SNVs; following electroporation into embryonic chick wings, the patient-*LARM2* sequence lacked activity (n=3; compare with the common *hLARM2* sequence).

Also, in a sporadic NPS case without ocular or renal involvement, we identified loss of heterozygosity of the *LARM* region. The *LARM*2 region was homozygous for four single nucleotide variations (SNVs) that were uncommon (each with an incidence of 16%) and one rare SNV (0.08%) (Fig. 5O). SNV-array showed several large homozygosity regions on chromosomes 9 (comprising *LMX1B*) and 17 (Fig. 5O), arguing for distant consanguinity in this individual. Functional analyses of the *LARM2* containing the five SNVs in the chicken bioassay showed that this *LARM2* variant was inactive (Fig. 5P). These results suggest that homozygosity for this rare haplotype is responsible for the autosomal recessive NPS, since the heterozygous carrier parents are unaffected.

## Discussion

In this report, we characterize two conserved *<u>L</u>mx1b*-<u>a</u>ssociated *cis*-<u>r</u>egulatory <u>m</u>odules (*LARM1* and *LARM2*) that are bound by Lmx1b and required to amplify *Lmx1b* expression in the limb to levels sufficient to accomplish limb dorsalization. Thus, *LARM1* and *LARM2* are two limb-specific *Lmx1b* enhancers responsible for establishing the correct dorso-ventral pattern.

Consistent with being limb-specific enhancers, *LARM1/2* mouse mutants do not carry other *Lmx1b*-associated abnormalities that might jeopardize survival, thereby offering an extraordinary opportunity to study the functional capacities of ventral-ventral limbs. Interestingly, *LARM1/2* mutants cannot walk but rather crawl because their limbs are unable to push-up their body. Indeed, the movement/locomotion of *LARM1/2* mutant limbs is reminiscent of that of fins. The LARM region is conserved through *Xenopus*, but not in fishes and raises speculation that the fin to limb transition may have required the acquisition of distal dorso-ventral asymmetries.

We also show that disruption of these enhancers can cause human pathology since loss-of-function variations in the *LARM* region are responsible for a limb-specific form of human NPS. This limited form of NPS is not associated with the typical risk of chronic kidney disease or glaucoma. Moreover, there is no protein-coding variation. Recognition that disruption of the *LARM* region can cause this limited form of NPS provides these patients with a more accurate assessment of their condition.

The NPS phenotype is attributed to haploinsufficiency, i.e, reduced levels of LMX1B. Our studies further characterize the pathogenicity of NPS to reduced levels of LMX1B. In one NPS family a single allelic deletion of LARM2 yields limb features diagnostic of NPS (incomplete limb dorsalization) indicating that LMX1B protein levels are below the normal patterning threshold. In another family, homozygosity of a functionally impaired LARM2 allele also yields limb features diagnostic of NPS. In both of these families, the remaining LARM1 enhancers, which demonstrate clear activity in mouse transgenesis and chicken assays, appear able to support LMX1B amplification to partially dorsalize the limb and avert a more severe ventral-ventral phenotype. Together our results point to the contribution of both allelic sets of LARM1/2 to get a fully functional dose of LMX1B.

Congenital limb anomalies are relatively common^14,15^ with syndromic forms associated with more than a hundred genes. The association of multiple affected organs (developmental pleiotropy) provides a clue to the affected gene and permits a high diagnostic yield. However, more than half of limb anomalies are isolated without other malformations, and the diagnostic yield of genetic evaluation remains low in these cases due, at least in part, to an emphasis on evaluating coding sequences. During morphogenesis, tissue-specific CRMs cause developmental pleiotropy by regulating genes in key developmental pathways in precise temporal and spatial patterns. Thus, tissue-specific CRMs are potential candidates to explain isolated limb anomalies. Our findings, as well as others linking limb-specific CRMs to limb anomalies^16–19^, support this concept. Characterization of CRM-disease associations represents a forthcoming opportunity in clinical genetics, not only for limb anomalies, but also for other isolated malformations.

## Methods

### Animal procedures

All animal procedures were reviewed and approved by the Loma Linda University Institutional Animal Care Use Committee (IACUC) or by the Bioethics Committee of the University of Cantabria and performed according to the EU regulations, animal welfare and 3R principles.

### Functional enhancer validation in chicken bioassays

We used a thymidine kinase (*tk*) promoter-driven GFP reporter (kind gift of Masanori Uchikawa)^24^ to generate enhancer constructs. Functional analyses of the *LARM1* and *LARM2* constructs were performed by electroporation into presumptive limb mesoderm of Hamburger and Hamilton stage (HH) 14 chicken embryos. Co-electroporation of a β-actin promoter-driven RFP construct (pCAGGS-RFP, kind gift from Cheryl Tickle)^25^ was used to determine transfection efficiency. Electroporation was performed using the CUY21 electroporation station (Protech International, Boerne, TX). Embryos were incubated for 48h before harvesting for visualization of GFP activity and digital image acquisition (Sony DKC-5000). To demonstrate that LMX1B could induce construct activity, pCDNA3.1-hLMX1B (kind gift from Roy Morello)^5^ was co-electroporated with p*tk*-LARM2-GFP into the ventral mesoderm of stage HH23 chicken limb buds.

### Cloning and site-directed mutagenesis

Primers used for the isolation of enhancer sequences from genomic DNA are listed in Table S1. Disruption of the Lmxb1 binding sites was performed using the QuikChange Lightning Site-Directed Mutagenesis Kit (Agilent Technologies, Santa Clara, CA) following manufacturer recommendations and confirmed by Sanger sequencing.

### In vivo transgenic reporter assays

Lmx1b associated regulatory modules were isolated from human genomic DNA and cloned into the hsp68-LacZ kindly provided by Dr. Ahituv^26^ with the primers listed in Table S1. The constructs were used to generate transgenic embryos (Cyagen transgenic service, Santa Clara, California). The embryos were harvested at e12.5 and processed for detection of LacZ activity.

### Analyses of published data

Limb ChIP-seq data were obtained from the Gene Expression Omnibus database (GEO, http://www.ncbi.nlm.nih.gov/geo/) under the accession numbers GSE84064 for Lmx1b, GSE42413 for H3K27Ac, GSE13845 for p300 and GSE42237 for both H3K27me and H3K4me. RNA Pol II and Med12 ChIP-seq data were available from Berlivet and coworkers^23^. Previously published data containing the genomic coordinates of interest were uploaded to the UCSC genome browser and converted to the mouse build mm10 using the liftover tool. Capture-C data were accessible under the GEO GSM2251518.

### CRISPR-Cas9 mediated enhancer knockout mice generation

The knockout mouse strain for the limb specific *Lmx1b*-associated regulatory modules (*LARM1/2*) was generated with the use of CRISPR-Cas9. Single guides RNAs (sgRNA) flanking the *LARM* locus listed in Table S1 were designed using breaking-Cas^27^. Cleavage efficiency of the selected sgRNAs was tested using cel-1 assay in mouse neuroblastema (N2a) cells (SIGMA) with the primer pairs listed in Table S1. Generation of knockout mice was performed following cytoplasmic injection of 50 ng/μl each sgRNA and 100 ng/μl of Cas9 mRNA, in injection buffer (1 mM Tris–HCl pH 7.5; 0.1 mM EDTA pH 7.5), into B6CBA mouse strain single cell embryos at the National Center of Biotechnology′s transgenic core facility (CNB-CSIC). Genotyping of founder (F0) mice for the identification of the desired deletion was performed with the use of the Phusion high fidelity polymerase (Thermo Scientific) with the primers listed in Table S1. A founder female B6CBA was bred to C57/BL6 for the generation of the *LARM1/2* knockout mice strain after performing Sanger sequencing of the PCR amplicons that confirmed a 7.564 bp deletion encompassing the *LARM* locus (Chr9 :33555354-33563008; Mouse July 2007(NCBI/mm9) assembly). The genotyping strategy included Sanger sequencing of the PCR amplicons for the off-target regions providing the highest score for each of the sgRNAs according to Breaking-Cas with the use of the primers listed in Table S1 confirmed no genetic alteration.

### MicroCT

Three-week-old mouse limbs were scanned with Skyscan1172 at 40 kV, 100 μA, and 27.03 μm pixel resolution. Subsequent reconstruction was performed using the N record reconstruction software.

### Skeletal preparations

After removal of the skin and viscera, animals were fixed in 95% ethanol. Alizarin red and alcian blue skeletal staining was performed following standard procedures.

### Histology

Three-weeks-old animals were subjected to intravascular perfusion of 4%PFA with the use of a peristaltic bomb. Gross morphologic and histologic analyses were performed on the limbs, skull, brainstem, kidneys and eyes. The soft tissues (brain, kidneys and eyes) were fixed in 10% phosphate-buffered formalin, while the limbs were decalcified then post-fixed with 10% phosphate-buffered formalin. The tissues were paraffin embedded following standard procedures and stained with hematoxylin and eosin.

### RNA isolation and quantification by real-time PCR

Fore- and hind-limb buds from e12.5 wild type and *LARM1/2* KO embryos were dissected in cold RNAse-free PBS. Fore- and hind-limb buds of individual embryos were pooled and total RNA was isolated using the RNeasy Mini Kit (Qiagen) for subsequent reverse transcription as per Superscript III (Life Technologies) manufacturer recommendations. Quantitative PCR (qPCR) was performed on the Mx3005P cycler using SYBRGreen PCR Master Mix (Life Technologies). Data were analyzed using the MxPro software (Agilent Technologies). Primers listed in Table S1 were manually designed to span an intron in order to prevent genomic DNA amplification. Relative *Lmx1b* expression levels in three individual biological replicates for each genotype were normalized to that of the s14 housekeeping gene. Mean values of *Lmx1b* expression levels were calculated, and for comparison between genotypes, the mean of *Lmx1b* expression in wild type was set to 1. Statistically significant differences were considered when *P*<0.01 (two-tailed, unpaired *t*-test).

### *In situ* hybridization

Whole-mount *in situ* hybridization was performed following standard procedures as previously described^28^ using a previously described antisense RNA probe for mouse *Lmx1b*^3^ and Human *LMX1B^5^.* To generate the chicken *LMX1B* probe *…*

### LARM screening in NPS patients

DNA from patients was extracted from blood according to standard methods. *LARM1* and *LARM2* were sequenced on an ABI Prism 3730XL Genetic Analyzer (Applied Biosystems, Courtaboeuf, France) using Big Dye Terminator v3.1 Cycle Sequencing Kit, after PCR amplification using the following primers pairs:

hLARM1_5p1-F GTGTAGGTTTGACGGTGGGATTTTCC,
hLARM1_5p2-R GGGATTTTCCTGGCATCAAACAGC,
hLARM1_3p1-F GCTGGAGCCCATGAGAAGATTGC,
hLARM1_3p2-R GACAGGGTTGGATTGGTCTCTTGG,
hLARM2_5p1-F CCCACGGCAGGAGTTATAAGCAAGG,
hLARM2_3p1-R CGGACCAGGAGAAACATTCTTCTGTG.

Copy number of *LARM1* and *LARM2* was studied by real time quantitative PCR using SYBR Green technology (Applied Biosystems®, Saint Aubin, France) with the following primers pairs: LARM1-F AATTAACGGCTCCTCCCTG, LARM1-R GCCTTCTTCCTACTTCTGTCA, LARM2-F GTCTCTGCCCCTCGCTGA, LARM2-R CGTGGGCAATATGGCTTTGAA. Quantification of the target sequences was normalized to an assay from *RPH Polymerase* (NR_002312), and the relative copy number was determined on the basis of the comparative ΔΔCt method using a normal control DNA as the calibrator.

## Acknowledgements

The authors are grateful to Dr Boris Keren for the analysis of the SNP array in family 2. This work was supported in part by grants from the Spanish Ministerio de Ciencia, Innovación y Universidades (M.A.R) (BFU2017-88265-P); the National Organization for Rare Disorders (K.C.O.) and the Loma Linda University Pathology Research Endowment Fund (K.C.O.).

## Author contributions

EH, FP, MAR and KCO conceived and designed the project. EH, CUP, CDS, LAI, ALG, and AN acquired and analyzed murine and human *LARM1/2* characterization studies in chickens. EH, CUP, ALG, and KCO acquired and analyzed human *LARM1/2* studies in transgenic mice. EH, KCO and MAR acquired and analyzed the CRISPR-cas9 knockout model mice. FP and SMH supervised, the acquisition and analysis of the human Nail-Patella Syndrome (NPS) data. FE, A-SJ, FF, J-MG, and CF acquired and analyzed human NPS data. EH, FP, KCO, MAR drafted the manuscript. All authors were involved in editing/revising the manuscript.

